# Atlantic and Indo-Pacific separation in *Palythoa* sibling species: phylogenomic analyses using ultraconserved elements

**DOI:** 10.64898/2026.04.26.720863

**Authors:** Lara Adele Jacobs Hansen, Maria E. A. Santos, Hiroki Kise, Nuba Zamora-Jordán, James Davis Reimer

## Abstract

The delineation of closely related species remains a persistent challenge in Zoantharia, where morphological plasticity and limited genetic differentiation complicate taxonomy. In this study, we investigated the phylogenetic relationship between the widely distributed sibling taxa *Palythoa tuberculosa* (Indo-Pacific) and *Palythoa caribaeorum* (Atlantic) using ultraconserved elements (UCEs) recovered from genome skimming. A dataset comprising 116 loci (35,699 bp) across 37 specimens from Brazil, the Red Sea, Okinawa, and New Caledonia was analysed using both concatenated maximum-likelihood and coalescent-based approaches. Phylogenetic reconstructions did not recover monophyletic relationships corresponding to either species or geographic origin, instead revealing intermixed lineages across the Indo-Pacific and Atlantic regions. Concordance factor analyses indicated low gene concordance and moderate site concordance, suggesting pervasive gene tree discordance rather than a lack of phylogenetic signal. These patterns are consistent with previous studies based on mitochondrial, nuclear, and reduced-representation datasets, indicating that increased marker resolution does not resolve species boundaries within this complex. The observed lack of differentiation may reflect ongoing or recent connectivity among populations, potentially facilitated by long-distance dispersal promoted by anthropogenic rafting or historical range expansion, biological invasion, or biological processes such as incomplete lineage sorting. The results support the hypothesis that *P. tuberculosa* and *P. caribaeorum* represent a species complex or a case of incipient speciation rather than fully distinct evolutionary lineages. These findings indicate that genome-scale data alone may be insufficient to resolve very recent divergences, supporting the need for integrative approaches to resolve complicated species boundaries in zoantharians.

## INTRODUCTION

Zoantharians (Cnidaria; Anthozoa; Zoantharia) occupy diverse marine habitats, including shallow tropical coral reefs, intertidal flats, and mesophotic environments, often colonising biotic substrates through overgrowth of other sessile invertebrates (Reimer and Fujii 2010; Carreiro-Silva et al. 2017; Reimer et al. 2021). Zooxanthellate zoantharians are among the most common shallow-water hexacorals and lack a calcareous skeleton, enabling them to outgrow reef-building corals and compete for space aggressively (Lustic et al. 2020). Particularly, species of the genus *Palythoa* have been increasingly associated with ecological phase shifts and reef degradation, where they can dominate benthic communities under conditions that limit the persistence of reef-building corals (Cruz et al. 2016; Reimer et al. 2021; Zamora-Jordán et al. 2026). As an ecologically and morphologically flexible genus, *Palythoa* spp. are therefore of growing interest in the context of anthropogenic environmental change.

Most *Palythoa* species incorporate sand grains and sediment into the coenenchyme and polyp body wall for structural support, environmental resilience, and predator defence (Haywick and Mueller 1997; Reimer et al. 2010; Santos et al. 2024). These features, alongside characters such as sphincter muscle anatomy and cnidome composition, have historically been used for species identification, however their taxonomic utility is often limited across broad spatial and environmental contexts (Low et al. 2016; Carreiro-Silva et al. 2017). This limitation is particularly evident in the widely distributed sibling pair *Palythoa tuberculosa* (Indo-Pacific) and *Palythoa caribaeorum* (Atlantic), which display many environmentally induced morphotypes. Factors such as sedimentation and seasonality can influence polyp density, diameter and thickness of colonies (Costa et al. 2012) while substrate type (e.g. volcanic, carbonate) can affect encrustation patterns and pigmentation (Reimer et al. 2006). Similarly, changes in irradiance can generate morphotypes easily mistaken for different taxa (Ong et al. 2013).

Consistent with these morphological challenges, previous molecular studies using sequences from both nuclear (internal transcribed spacer (ITS) region) and mitochondrial (cytochrome oxidase subunit I (COI) and 16S ribosomal DNA) markers have shown limited resolution among closely related *Palythoa* species, including this sibling pair (Sinniger et al. 2005; Reimer et al. 2007; Mizuyama et al. 2018; López et al. 2019). This lack of resolution is partly explained by the slower rate of mitochondrial evolution found in Anthozoa, where substitution rates are estimated to be 10 to 100 times lower than in other metazoans, with Zoantharia among the slowest evolving groups (Shearer et al. 2002; Huang et al. 2008; Quattrini et al. 2023).

Patterns of marine biodiversity are strongly shaped by biogeographic barriers that limit connectivity and generate allopatric speciation between ocean basins (Santos et al. 2026). In tropical marine systems, the separation between the Atlantic and Indo-Pacific ocean basins is largely maintained by the African landmass and the cold, upwelling-dominated Benguela Current on the southwest African margin, with unsuitable shallow-reef habitat around the Cape (Dudoit et al. 2018). In combination with the episodic nature of the Agulhas Current leakage, these factors are expected to restrict gene flow between the Indo-Pacific and Atlantic for warm-water taxa (Beal et al. 2011; Schleyer et al. 2018). This pattern has been widely documented across multiple marine groups (Schwaninger 2008; Stelbrink et al. 2010), but may be less prevalent in zoantharians, and some hexacorals (Arrigoni et al. 2018; Dudoit et al. 2021).

Recent phylogenomic analyses have further supported the lack of clear species boundaries between *P. caribaeorum* and *P. tuberculosa*, including evidence from reduced representation sequencing (eg. ezRAD), single nucleotide polymorphisms (SNPs) and transcriptomic loci (Dudoit et al. 2021). These findings indicate inconclusive species-level relationships as well as evidence for polyphyly in the sibling pair. The consistency of these results across marker types indicates that the relationship between this sibling pair remains a persistent and unresolved feature in *Palythoa* systematics, and confirms that the taxonomic complexity in this group is biological rather than a limitation of marker choice or methodology.

Here, we build on these findings by applying ultraconserved elements (UCE) phylogenomics to investigate the relationship between *P. caribaeorum* and *P. tuberculosa* across Atlantic and Indo-Pacific distributions. The present study includes geographic sampling across four global locations: the Red Sea, Okinawa and New Caledonia in the Indo-Pacific, and Brazil in the Atlantic, and represents the first application of UCEs to this sibling pair. Specifically, we assess genetic differentiation between these taxa and test whether these lineages exhibit biogeographic segregation or instead reinforce previously observed patterns of unclear species boundaries, using both concatenated maximum likelihood and coalescent-based species tree approaches across many independent UCE loci.

UCEs have emerged as a powerful tool for addressing taxonomic uncertainty in Anthozoans, providing thousands of orthologous loci across taxa and improving resolution across various taxonomic levels from recent speciation to deep-time order and subphyla levels (Faircloth et al. 2012; Quattrini et al. 2018; Erickson et al. 2021; McFadden et al. 2021). Their conserved core regions enable reliable alignment and locus recovery across species, and more variable flanking regions retain informative genetic variation, making them well suited for resolving relationships characterised by weak genetic differentiation or recent divergence, as observed in *Palythoa*. Given the ecological importance and widespread distribution of *Palythoa*, clarifying evolutionary relationships within this genus is essential for understanding patterns of biodiversity, connectivity, and species boundaries in zoantharians.

## MATERIAL AND METHODS

### Specimen sampling

Specimens of *P. tuberculosa* (Esper, 1791) from the Red Sea (n = 10) and New Caledonia (n = 10), as well as *P. caribaeorum* (Duchassaing de Fonbressin & Michelotti, 1860) from Brazil (n = 10) were provided by the MISE Lab (University of the Ryukyus). Additional *P. tuberculosa* specimens were collected (n = 12) using SCUBA diving from Ginowan (26°17’23.4”N 127°44’46.4”E), Mizugama (26°21’32.9”N 127°44’18.1”E), Oku Beach (26°51’14.2”N 128°16’59.4”E), and Adan Beach (26°49’29.6”N 128°18’45.5”E) in Okinawa Island, Japan, in April 2025, with morphology shown in **Figure 1**.

**Figure 1.**
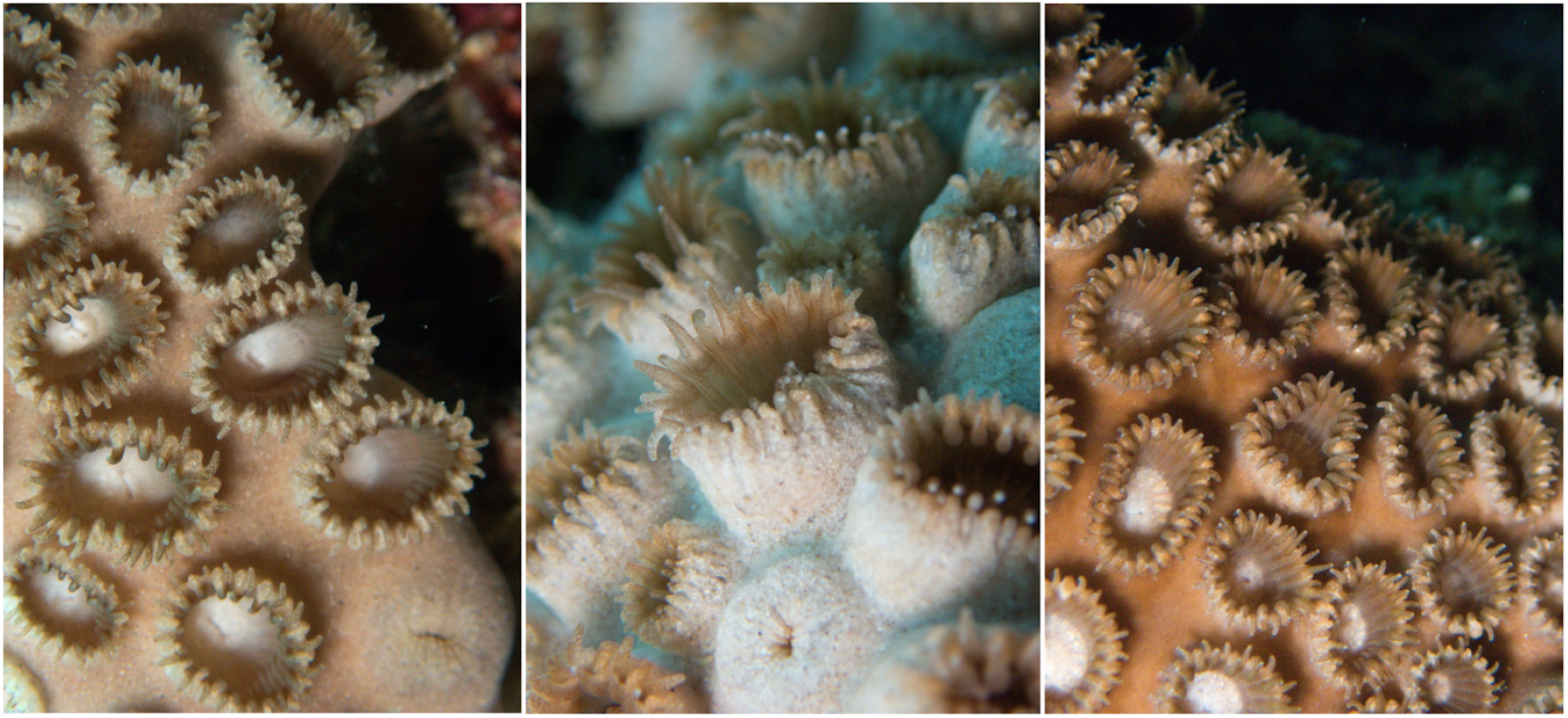
Images of *in situ Palythoa tuberculosa* colonies taken during the time of sampling in Ginowan, Okinawa.

At least ten polyps were collected per colony, and colonies were sampled at least two metres apart to avoid pseudoreplication. Samples were then stored at room temperature in 99.5% ethanol prior to DNA extraction. Sampling locations are shown in **Figure 2**.

**Figure 2.**
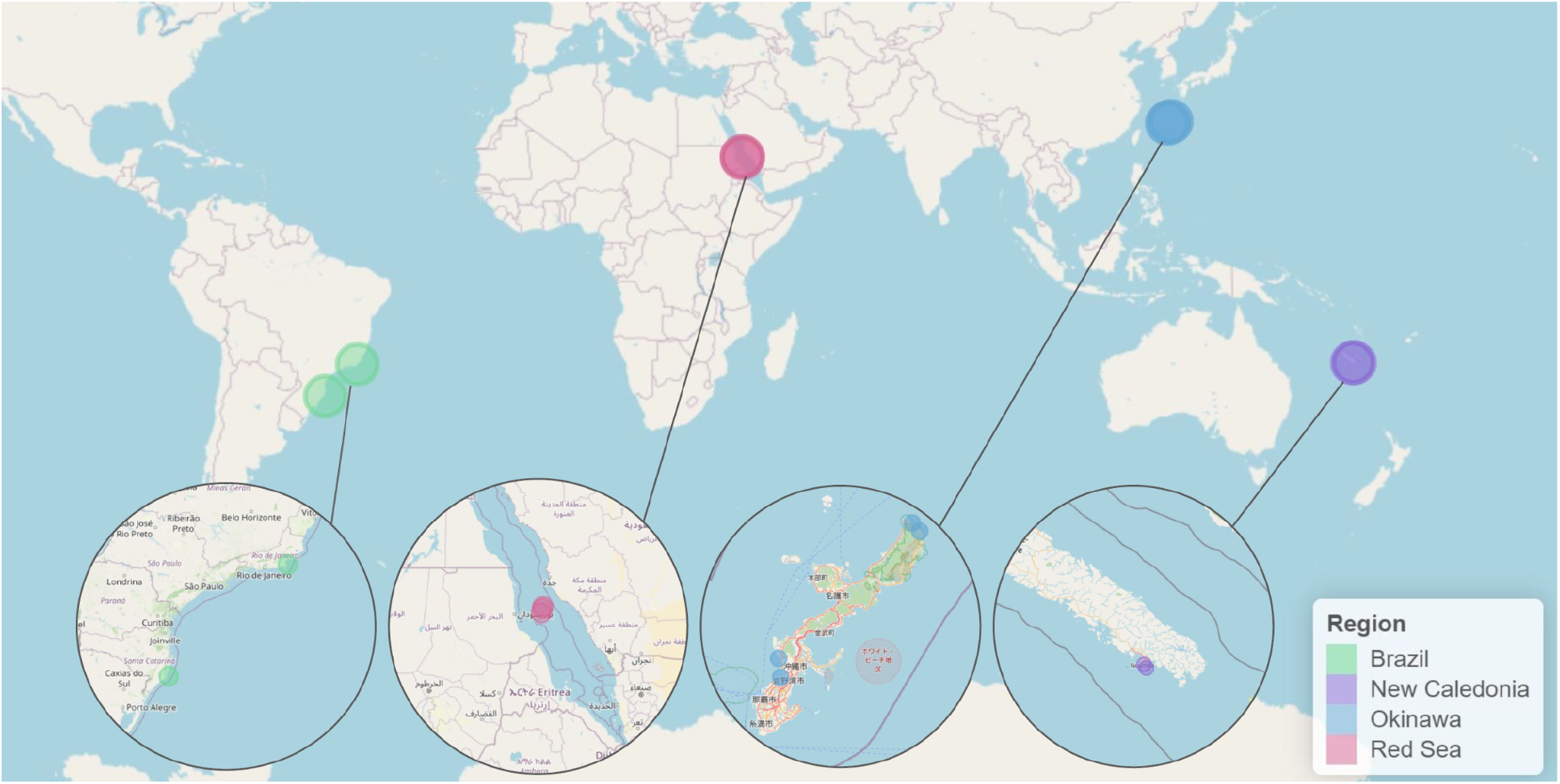
Geographical locations of the complete specimen list sent to the National Institute of Advanced Industrial Science and Technology AIST, including inlets of each region to show locations at a higher resolution. Colours used for each region are homogenised across all tables and figures (Brazil: green, New Caledonia: purple, Okinawa: blue, Red Sea: red).

### Molecular laboratory processing

At least two days after collection, genomic DNA was extracted using either the DNeasy Blood and Tissue Kit (Qiagen) or Monarch Genomic DNA Purification Kit (NEB, T3010), following manufacturer protocols, and eluted in Tris–EDTA buffer. DNA concentrations were measured with a NanoDrop spectrophotometer, then stored at -20 °C; only samples with a concentration of at least 10 ng µL^-1^ of DNA in at least 20 µL were retained for further analyses.

Library preparation for all samples was carried out using NEBNext Multiplex Oligos adaptor kit at the National Institute of Advanced Industrial Science and Technology (AIST, Tsukuba, Japan). DNA concentration after library preparation was quantified using a Qubit dsDNA HS assay kit with a Qubit 4.0 Fluorometer (Thermo Fisher Scientific, Waltham, USA) and ranged from 0.20 to 63.6 ng of total DNA; with a mean of 36.06 ± 19.75 (SD). Samples with insufficient DNA quality or concentration were excluded. Libraries were enriched with ten PCR cycles and sequenced on Illumina NovaSeq X Plus with paired-end sequencing (150 base pairs). Genome skimming was used to generate low-coverage genomic data for subsequent UCE recovery.

Raw sequencing reads generated in this study have been deposited under BioProject accession PRJNA1450055, with associated metadata including collection site and depth. Ten additional zoantharian taxa that were used as outgroups were obtained from publicly available sequencing datasets by Quattrini et al. (2018) and retrieved from the NCBI Sequence Read Archive (SRA) under BioProject accession PRJNA588468.

### UCE assembly and matrix construction

All raw sequence files, including outgroups, were processed using PHYLUCE v1.6.8 (Faircloth et al. 2012; Faircloth 2016) and corresponding standard online tutorials. Quality trimming of raw FASTQ reads was conducted through Trimmomatic v0.39 (Bolger et al. 2014) and assembled de novo using SPAdes v3.12.0 (Prjibelski et al. 2020).

UCE loci were then identified from assembled contigs using the probe set *Cowman_etal_APPENDIX_C-hexa-v2-final-probes*.*fasta* (Cowman et al. 2020), applying minimum identity and coverage thresholds of 80%, consistent with previous methods (Faircloth et al. 2012). A total of 2,476 UCE loci were recovered. Corresponding UCE regions were extracted from assembled contigs and compiled into locus-specific alignments.

Multiple sequence alignments were performed using MAFFT v7.525 (Katoh and Standley 2013), with loci initially filtered to retain alignments of at least 100 base pairs (bp), with representation in at least 65% of taxa, and with a maximum in-locus divergence of 0.2. After multiple sequence alignment, TrimAl v1.4 (Capella-Gutiérrez et al. 2009) was used to prune poorly aligned and ambiguous regions with automated parameters. This resulted in 1,534 trimmed alignments (mean length of 257 bp, total alignment length of 394,246 bp).

Final per-locus datasets were created by further filtering alignments based on minimum taxon occupancy thresholds of 45%, 55%, 65%, and 70%. Different taxon-occupancy thresholds were considered to assess robustness of missing data. As higher thresholds retained substantially fewer loci (**Supplementary Table S1**) samples with extremely low UCE recovery (<200 loci) were excluded to improve matrix completeness. The final phylogenomic dataset was generated using a 55% occupancy matrix comprising 116 loci (35,699 bp; 6,234/17.5% informative sites) producing a concatenated PHYLIP alignment and a per-locus partition file. A summary of the final matrix summary is shown in **Table 1**.

**Table 1.**
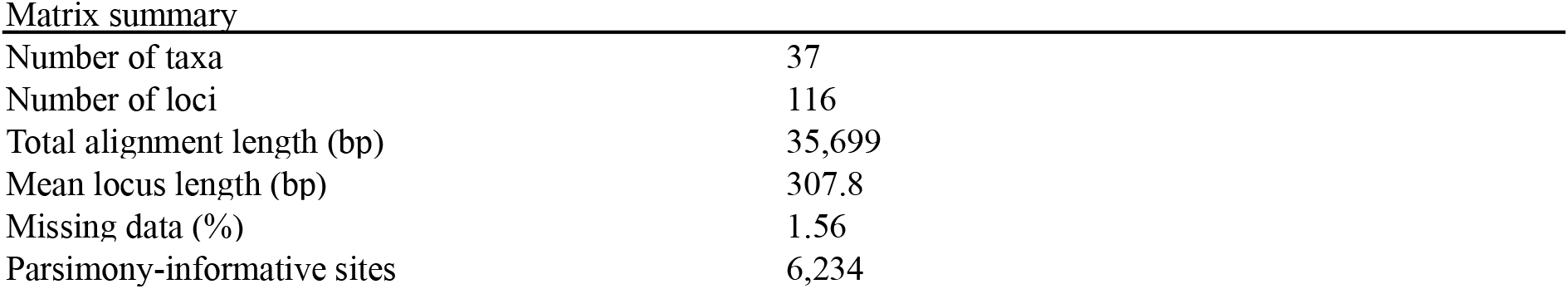
Summary statistics for the final 55% occupancy UCE dataset used in phylogenetic analyses.

### Phylogenetic analyses

Maximum-likelihood (ML) phylogenies were inferred with IQ-TREE v3.0.1 (Nguyen et al. 2015; Minh et al. 2020). Analyses were performed using a per-locus partitioning scheme (Chernomor et al. 2016) with automatic substitution model selection for each partition using ModelFinder (Kalyaanamoorthy et al. 2017) under the Bayesian Information Criterion (BIC). Branch support was estimated with ultrafast bootstrap approximation (UFBoot2) with 1,000 replicates (Hoang et al. 2018).

Gene and site concordance factors were calculated in IQ-TREE to quantify concordance among loci and alignment sites across the inferred phylogeny. Gene concordance factors (gCF) represent the proportion of gene trees supporting a given branch of the reference tree, whereas site concordance factors (sCF) represent the proportion of informative alignment sites supporting that branch. Concordance factors were calculated by comparing individual locus trees against the ML tree inferred from the concatenated UCE alignment.

Species relationships were additionally inferred using ASTRAL v5.7.8, which uses the ASTRAL-III algorithm under the multispecies coalescent framework (Mirarab et al. 2014; Zhang et al. 2018). Gene trees were first estimated for individual UCE loci and used as input for ASTRAL, which reconstructs species trees by maximizing quartet agreement across gene trees. Node support was assessed using local posterior probabilities (LPP), which estimate the probability that a given branch is correct based on quartet frequencies observed across gene trees. To reduce noise from sparsely sampled loci, gene-tree analyses retained loci present in at least 12 taxa, to balance locus coverage and taxon representation. Topological similarity among phylogenetic trees was evaluated using the normalised Robinson–Foulds (RF) distance.

All trees were rooted using zoantharian taxa from the genera *Parazoanthus, Epizoanthus, Hydrozoanthus, Nanozoanthus, Microzoanthus*, and *Zoanthus* and visualised using Interactive Tree of Life (iTOL) (Letunic and Bork 2024). All bioinformatic computations were executed on the NIG Supercomputer at the Research Organisation of Information and Systems (ROIS) National Institute of Genetics of Mishima, Shizuoka, Japan.

## RESULTS

The final UCE matrix at a 55% taxon-occupancy threshold comprised 116 loci across 37 taxa, including *P. caribaeorum* from Brazil (n = 5), *P. tuberculosa* from the Red Sea (n = 8), New Caledonia (n = 9), and Okinawa (n = 5), with a total concatenated alignment length of 35,699 bp and a mean locus length of 307.8 bp. The dataset contained 6,234 parsimony-informative sites. Higher taxon occupancy thresholds, which retained substantially fewer loci (65% occupancy: 23 loci; 70% occupancy: 13 loci), were not considered further.

The concatenated ML phylogeny **(Figure 3)** was broadly consistent with the species relationships inferred using ASTRAL **(Figure 4)**. Both approaches recovered a clade containing intermixed *P. caribaeorum* and *P. tuberculosa* samples, with moderate support (UFBoot median = 74.5; LPP median = 0.60). Within the studied *Palythoa* sibling species, samples from Okinawa, the Red Sea, and New Caledonia did not cluster together by geographic region, and internal branches within the ingroup were short, resulting in a shallow topology. This pattern is illustrated in the panel containing only *Palythoa* (**Figure 3B**), where the branch separating *P. heliodiscus* from the focal cluster is longer than the branches within the sibling pair. As in the ML analysis, sequences from specimens of *P. caribaeorum* did not form a reciprocally monophyletic clade relative to *P. tuberculosa*, and were scattered among *Palythoa* lineages from the Indo-Pacific. Nevertheless, topological similarity between the ML and ASTRAL trees was moderate (normalised RF distance = 0.735). To assess the effect of gene-tree sampling on species-tree inference, loci were filtered to retain gene trees containing at least 12 taxa and, in a stricter analysis, 15 taxa. Both ASTRAL trees showed high topological similarity (normalised RF distance = 0.588), indicating that the inferred relationships were largely consistent across sampling thresholds.

**Figure 3.**
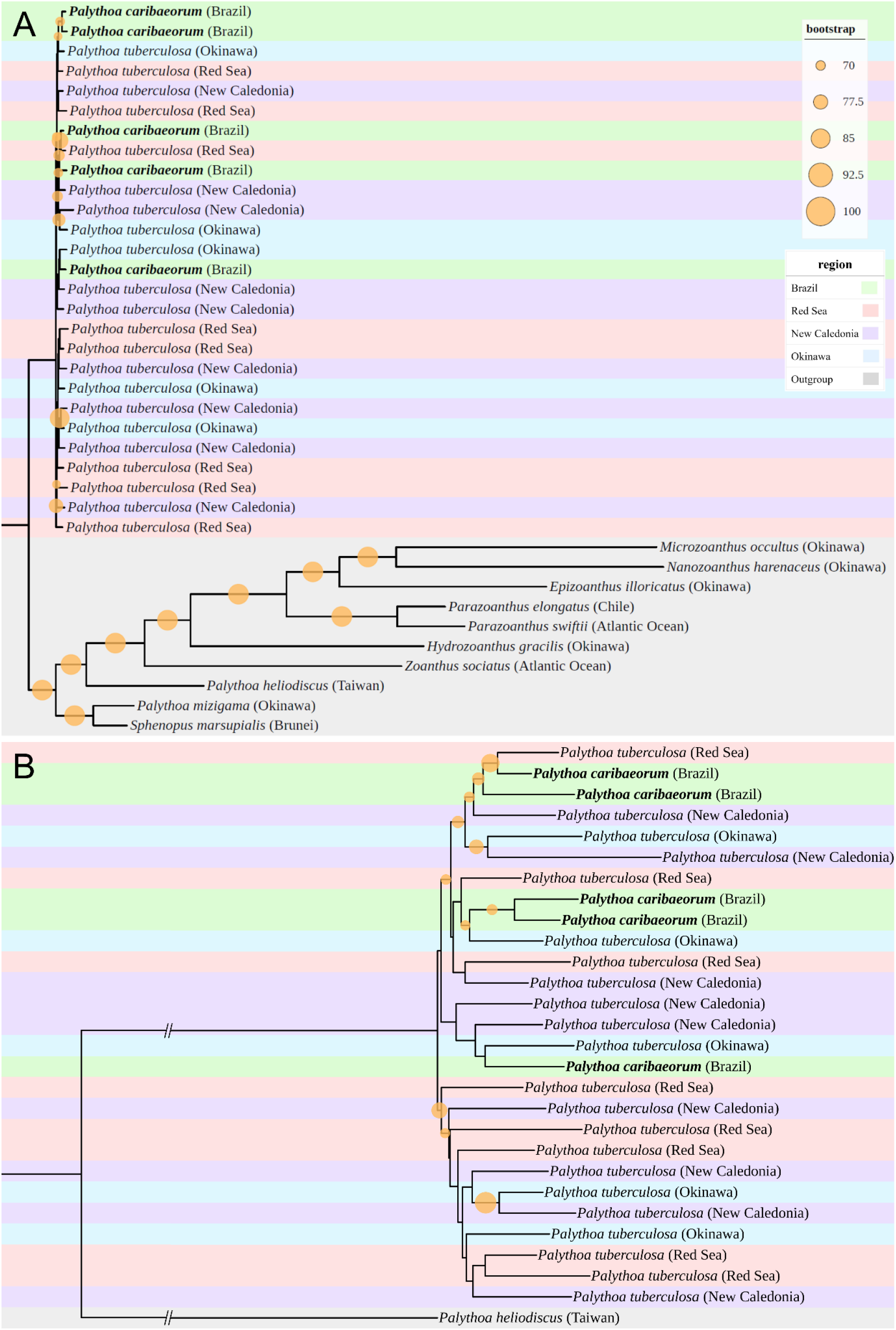
Maximum likelihood phylogeny reconstructed from the concatenated UCE alignment at a 55% occupancy threshold, using IQ-TREE. Node support is shown as ultrafast bootstrap values, represented by circles scaled by support value. (**A**) Full tree including outgroup taxa. **(B)** Inset showing the *Palythoa* ingroup with a single outgroup (*P. heliodiscus*) for visual clarity. Tip labels are colour-coded by sampling region and outgroups are shown in grey. For visualisation purposes, the basal branch separating the outgroups from the ingroup was truncated, while all other branch lengths are proportional to inferred substitutions.

**Figure 4.**
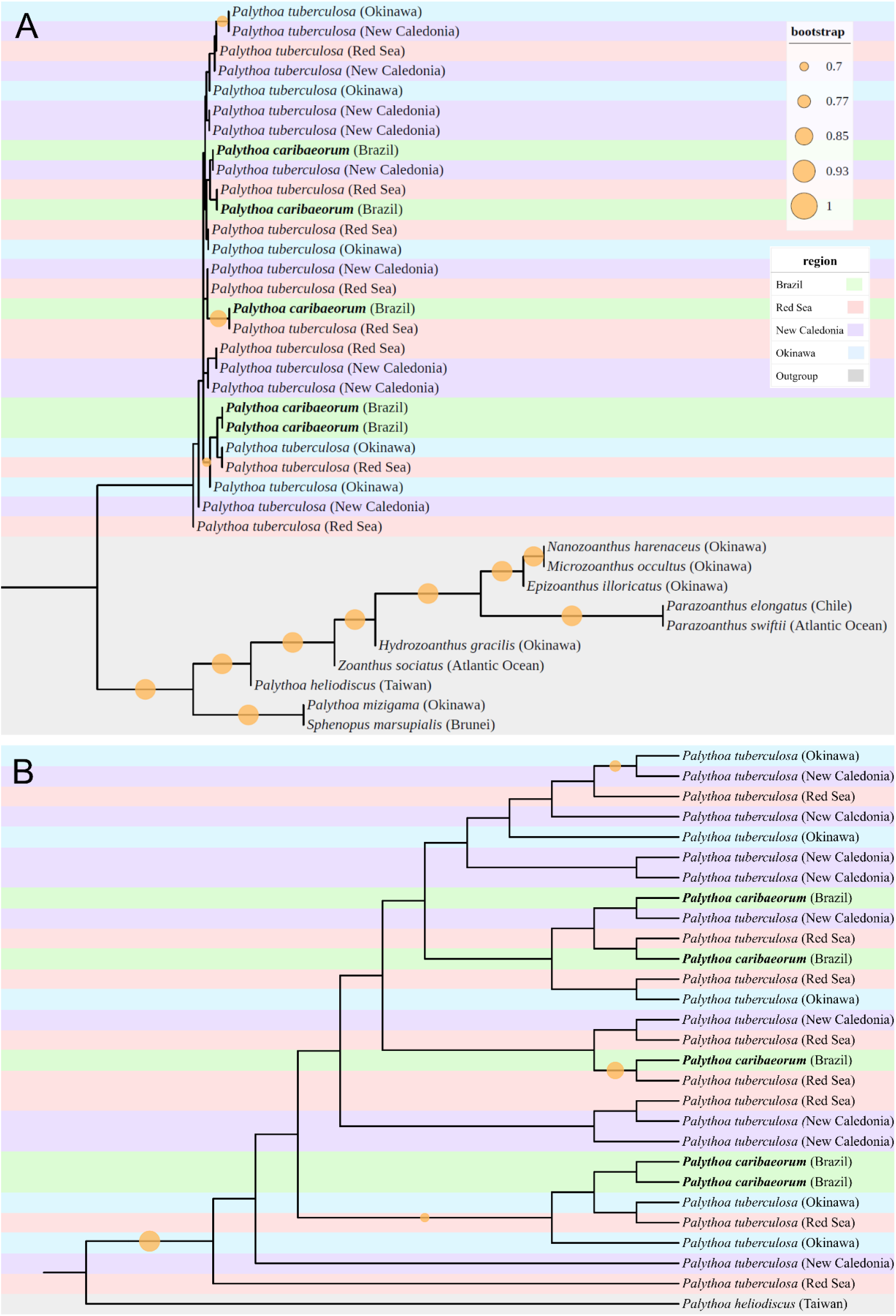
ASTRAL species tree inferred from individual UCE gene-trees (55% occupancy threshold, 116 UCE loci). **(A)** Full species tree including outgroup taxa. **(B)** Inset showing the *Palythoa* ingroup with a single outgroup (*P. heliodiscus*) for visual clarity. Node support is shown as local posterior probabilities (LPP), shown by circles scaled by support value. Tip labels are colour-coded by sampling region and outgroups are shown in grey.

Although concatenated ML analyses recover moderate to high bootstrap support (median UFBoot = 74.5), gCF were low across most internal branches (median gCF = 3.8%, mean = 18.6%, range = 0-83.5%) (**Figure 5A**). In contrast, sCF were moderate overall (median sCF = 40.3%, mean = 43.2%, range = 27.0-91.2%) (**Figure 5B**). Comparison of gCF and sCF across internal nodes revealed that site-level support generally exceeded gene-tree concordance (**Figure 6B**). Most branches showed low gCF values but moderate sCF values, with only a small number of nodes approaching similar values for both metrics. Most nodes fell above the diagonal line (representing the 1:1 relationship between gCF and sCF), showing that sCF were typically higher than gCF across internal branches of the tree (**Figure 5B**).

**Figure 5.**
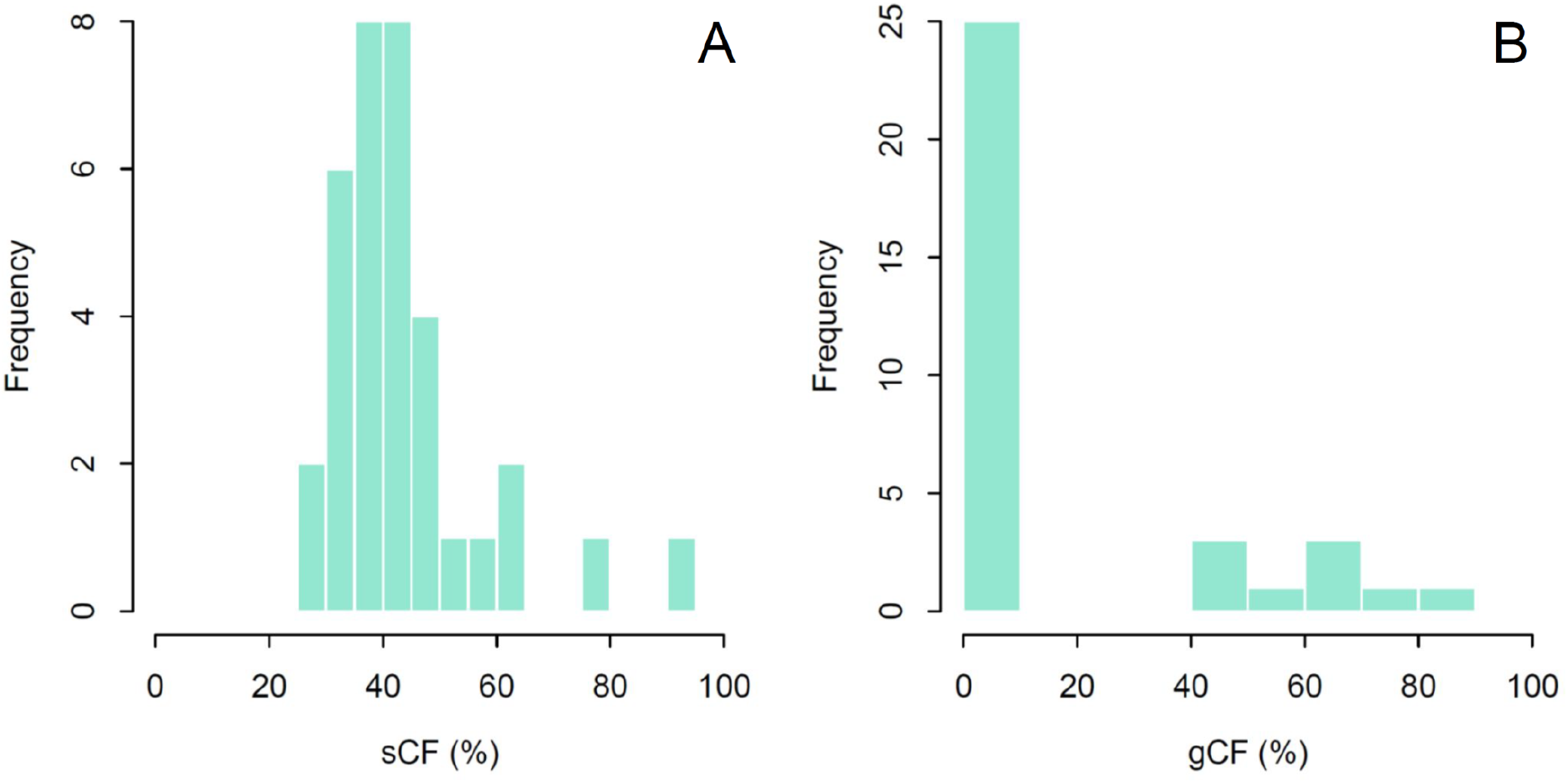
Distribution of concordance factors across nodes of the phylogeny. Each value corresponds to one internal node in the tree. Site concordance factors (sCF) measure how many informative sites in the alignment support that branch (Panel A). Gene concordance factors (gCF) measure how many gene trees support a given branch (Panel B).

**Figure 6.**
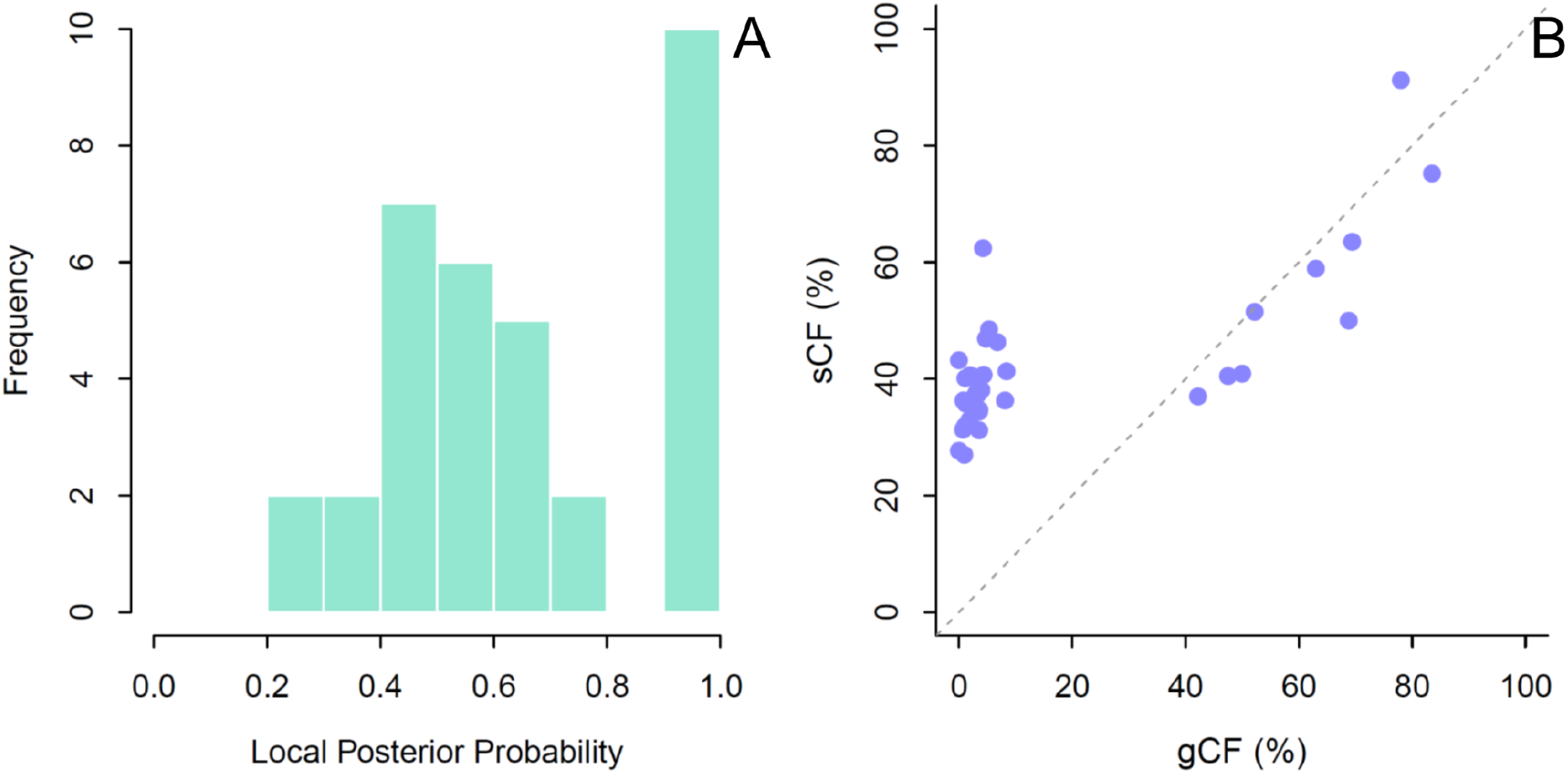
Summary of support across the tree. The histogram displays the distribution of local posterior probability (LPP) values across internal nodes of the ASTRAL species tree (Panel A). The scatter plot compares two concordance metrics where each point is an internal node in the phylogeny, with a dashed diagonal line indicating where gCF = sCF (Panel B).

The distribution of node support values in the ASTRAL species tree showed a range of local posterior probabilities (LPP) across internal branches (**Figure 6A**). Support values ranged from 0.20 to 1.00, with most values within intermediate support levels (median = 0.60; mean = 0.66). Approximately 73.5% of internal nodes exceeded 0.5, whereas fewer nodes reached higher support thresholds (29.4% ≥ 0.8; 26.5% ≥ 0.95). No internal node within the focal sibling-pair cluster reached LPP ≥ 0.95; the highest support was observed for a Brazil/Red Sea pair (LPP = 0.92). Node support summaries for all analyses are shown in **Supplementary Table S2**.

## DISCUSSION

The phylogenomic analyses using ultraconserved elements presented in this study provided no evidence of resolving species boundaries between Indo-Pacific *Palythoa tuberculosa* and Atlantic *Palythoa caribaeorum*, suggesting a species complex. Both the concatenated ML analysis (**Figure 3**) and the coalescent-based species tree (**Figure 4**) recover similar topologies, indicating overall consistency across methods. However, the RF distance indicates partial disagreement between reconstructions, while preserving the same broad phylogenetic structure. The lack of independently forming clades among sibling species with no geographical clustering may indicate either a very recent divergence event (incipient speciation) or maintained gene flow between lineages (Degnan and Rosenberg 2009). While moderate bootstrap support was observed across several internal nodes in the concatenated tree, concordance factor analyses showed that these relationships were supported by only a small proportion of gene trees (**Figure 5**), indicating substantial gene tree heterogeneity among UCE loci. Further, moderate sCF values suggest that phylogenetic signal was present within the alignment, although individual loci often supported alternative topologies. Additionally, the ≥12 taxon threshold in the ASTRAL species tree allowed inclusion of a greater number of loci while producing topologies consistent with those inferred under stricter filtering, suggesting that increased locus sampling did not compromise species tree inference despite moderate levels of missing data.

The short internal branches observed within the ingroup contrast with the deeper divergence separating the sibling cluster from the outgroup (*P. heliodiscus*), a pattern associated with low phylogenetic resolution in anthozoans (Quattrini et al. 2023). Short internodes reflect limited evolutionary time between divergence events, resulting in fewer informative sites to distinguish evolutionary lineages more generally (Degnan and Rosenberg 2009; Quattrini et al. 2023). Under these conditions, incomplete lineage sorting is a plausible underlying mechanism, as ancestral polymorphisms can persist across divergence events and fail to coalesce within the time interval between successive speciation events (Maddison 1997; Degnan and Rosenberg 2009). This leads to substantial gene tree discordance across the genome, consistent with the low gene concordance factors observed here. Introgression and hybridisation represent alternative processes that can also generate gene tree discordance (Heled and Drummond 2010). Although concatenated analyses can yield moderate bootstrap support by averaging across loci, such support does not necessarily reflect agreement among gene trees and may obscure underlying conflict (Kubatko and Degnan 2007; Whitfield and Lockhart 2007). Similar patterns have been widely reported in phylogenomic studies, where genome-scale datasets often fail to resolve recent divergences due to short internodes (Giarla and Esselstyn 2015). This pattern is particularly common in lineages with a large effective population size (N_e_) (Tan et al. 2023). While N_e_ of *P. tuberculosa* and *P. caribaeorum* have not been directly estimated, several life-history traits, including high reproductive output, extensive clonal growth, broadcast spawning, and high dispersal capacity suggest that large N_e_ may be maintained (Boscolo and Silveira 2005; Hirose et al. 2011; Polak et al. 2011; Santos and Reimer 2018), which would further prolong the persistence of incomplete lineage sorting times.

The persistence of this unresolved phylogenetic pattern among these two *Palythoa* sibling species (as well as in some other *Zoanthus* and *Palythoa* spp.) has also been documented in previous molecular studies using mitochondrial markers (COI, 16S rDNA) and the nuclear ITS region, reporting low sequence divergence and limited phylogenetic resolution (Sinniger et al. 2005; Mizuyama et al. 2018). For mitochondrial markers, these findings are expected given the slow rate of mitochondrial evolution in Anthozoa, which constrains the accumulation of phylogenetically informative variation at shallow timescales (Shearer et al. 2002). However, the similarly limited resolution observed in the nuclear regions is not attributable to mitochondrial rate constraints. Furthermore, this lack of resolution persists even when utilising higher-resolution nuclear datasets; reduced-representation genomic approaches, such as ezRAD and SNP-based analyses, have similarly recovered shallow divergences, weak population structure, and non-monophyletic species relationships among *Palythoa* sibling species (Dudoit et al. 2021). Consistently, the present UCE-based results extend this pattern by demonstrating that even many nuclear loci fail to recover consistent species boundaries within the *P. tuberculosa* and *P. caribaeorum* complex, therefore, further proving that the unresolved phylogenetic relationships between the pair reflect underlying biological processes rather than marker limitations (Dudoit et al. 2021). That being said, genome skimming approaches, while efficient, typically yield lower per-locus coverage than target capture and may therefore provide reduced resolution at shallow phylogenetic depths. In Anthozoa, UCE datasets derived from genome skimming can recover large number of loci, yet often place poorly supported splits near the tips of the tree, consistent with reduced phylogenetic information for recent divergences (Quattrini et al. 2023). Similar results have been reported across several anthozoan groups characterised by short internodes, including scleractinians and octocorals (Kitahara et al. 2016; Erickson et al. 2021). For example, in the scleractinian genus *Pocillopora*, genome-scale datasets have resolved deeper phylogenetic relationships but failed to distinguish among morphologically plastic species complexes (Oury et al. 2023). Likewise, mesophotic corals (e.g. *Leptoseris, Agaricia*) and reef-building taxa (e.g. *Acropora, Porites*) have shown discordant or weakly supported relationships at shallow phylogenetic levels (Rosser et al. 2017; Gijsbers et al. 2023). Together, these findings indicate that unresolved relationships among recently diverged lineages may be a common feature of anthozoan evolution.

On the other hand, the possibility of an invasive or recent range expansion should also be considered as a potential mechanism to partially explain the absence of the ingroup differentiation. Recent ecological and endosymbiont (Symbiodiniaceae) research supports the hypothesis that *P. caribaeorum* may be a recent coloniser in the Canary Islands derived from Indo-Pacific populations of *P. tuberculosa* (Zamora-Jordán et al. 2026). The phylogenetic results also showed Atlantic *P. caribaeorum* specimens to be nested within Indo-Pacific *P. tuberculosa*, and local Symbiodiniaceae communities being dominated by the Indo-Pacific generalist endosymbiont *Cladocopium proliferum*, displaying high levels of clonality and reduced allelic diversity. The lack of differentiation in endosymbiont communities suggests a potential founder effect following recent introduction, and vertical transmission of Symbiodinaceae profiles, further reinforcing the absence of clear biological boundaries between these sibling host taxa (Zamora-Jordán et al. 2026). In combination with the genomic results presented here, these findings support ongoing or recent connectivity among populations, driven by long-distance dispersal or range expansions.

Correspondingly, *Palythoa* spp. likely have potential for long-distance dispersal (Ryland et al. 2000; Polak et al. 2011; Dudoit et al. 2021) via both hermaphroditism and rafting, and exchanges between the Atlantic and Indian Oceans cannot be excluded. Broadcast spawning produces planktonic planulae, which may traverse thousands of kilometers under favourable circulation (Cowen and Sponaugle 2009). Episodic Agulhas leakage also pulses Indian Ocean water into the South Atlantic, creating rare opportunities for east to west transfer, with increased chances in exceptional years (Floeter et al. 2008; Cord et al. 2025). In addition, colony fragments or encrusted coenenchyme can raft on natural substrates (pumice, macroalgae, coral rubble) and anthropogenic flotsam, extending survival times well beyond documented larval durations (Polak et al. 2011; Santos and Reimer 2018). Such low-frequency, high-distance events would be insufficient to maintain routine gene flow across the Benguela barrier, but these systems, alongside human-mediated transport, provide plausible vectors for sporadic colonisation or biological invasion. Additionally, the effects of climate change and reduced macroalgal communities (Moreno-Borges et al 2022; Zamora-Jordán et al. 2024) may favour competitive colonisation behaviour of *P. caribaeorum* on benthic substrates (Lustic et al. 2020; Lambre et al. 2024).

In this context, incorporating population-level processes with a more comprehensive specimen dataset from the Atlantic Ocean is needed to characterise *P. caribaeorum* populations. Many coral groups including scleractinians, hydrocorals, and octocorals show lower species richness in the Atlantic due to simpler coastal geomorphology, reduced habitat heterogeneity, and westward direction of current flow (Ruiz-Ramos et al. 2014; Veron et al. 2015; McFadden et al. 2025). In contrast, the Indo-Australian Archipelago represents a major centre of coral diversification, where complex geography drives speciation, while oceanographic connectivity facilitates gene flow, resulting in weaker population structure within species (Bowen et al. 2016). Investigating whether similar patterns in population structure occur within *Palythoa* across ocean basins represents a valuable direction for future work.

This study represents the first application of UCEs to investigate the phylogenetic relationship between *P. tuberculosa* and *P. caribaeorum*, providing a genome-scale assessment. Overall, our results prove that these lineages do not conform to any biogeographic segregation, reinforcing previously observed patterns of unclear species boundaries. Although genome-scale approaches have transformed evolutionary clarity across Hexacorallia through universal and clade-specific probe sets (eg. Anthozoa-wide and Hexacorallia-focussed probe sets) (Faircloth et al. 2015; Quattrini et al. 2018; Cowman et al. 2020), their application within Zoantharia has been limited and only recently expanded through reduced-representation and UCE-based studies. However, because universal probe sets may be less informative, developing a zoantharian-specific UCE probe set may improve locus recovery and resolution over shallow evolutionary timescales. In addition, future work should integrate morphology, Symbiodinaceae profiles, or temporal calibration. In particular, divergence dating (molecular clock) would allow direct comparison with historical biogeographical events (Rieux and Balloux 2016) and re-evaluation of species boundaries, clarifying the interoceanic conspecificity observed between the pair. Quantifying genetic divergence using UCE-derived distances, analogous to COI-based approaches (Sinniger et al. 2010), and integrating these with multi-species coalescent methods would strengthen inferences about species boundaries under gene tree discordance. Resolving phylogenetic structure within *Palythoa* is essential to address persistent uncertainty, particularly in the context of ongoing environmental change. Further research is required to characterise the recent range expansion, biological invasion, and evolutionary processes in shaping the present distributions of this sibling pair. As *Palythoa* species are increasingly recognised as dominant components of benthic reef communities and altering community structure, taxonomic clarity is directly relevant for evaluating ecological function in response to environmental change and assessing biodiversity. Together, these considerations identify *Palythoa* as a priority for integrative taxonomy, combining genome-scale data with ecological context to resolve species boundaries.

## Ethical Approval

No animal testing was performed during this study.

## Sampling and Field Studies

All necessary permits for sampling and observational field studies have been obtained by the authors from the competent authorities. This study complies with the Convention on Biological Diversity (CBD) and the Nagoya Protocol.

## Data Availability

The raw sequencing reads that support the findings in this study have been deposited in the Sequence Read Archive under BioProject accession PRJNA1450055.

## Author Contribution Statement

LAJH: Sampling, data curation, formal bioinformatics analyses, data interpretation, main author.

MEAS: Sampling, investigation, data interpretation, conceptual input.

HK: Methodology, sequencing, bioinformatic support, conceptual input. NZJ: Conceptual input, manuscript revision.

JDR: Conceptualisation, investigation, funding acquisition, supervision, manuscript revision.

## SUPPLEMENTARY MATERIAL

**Supplementary Table 1.**
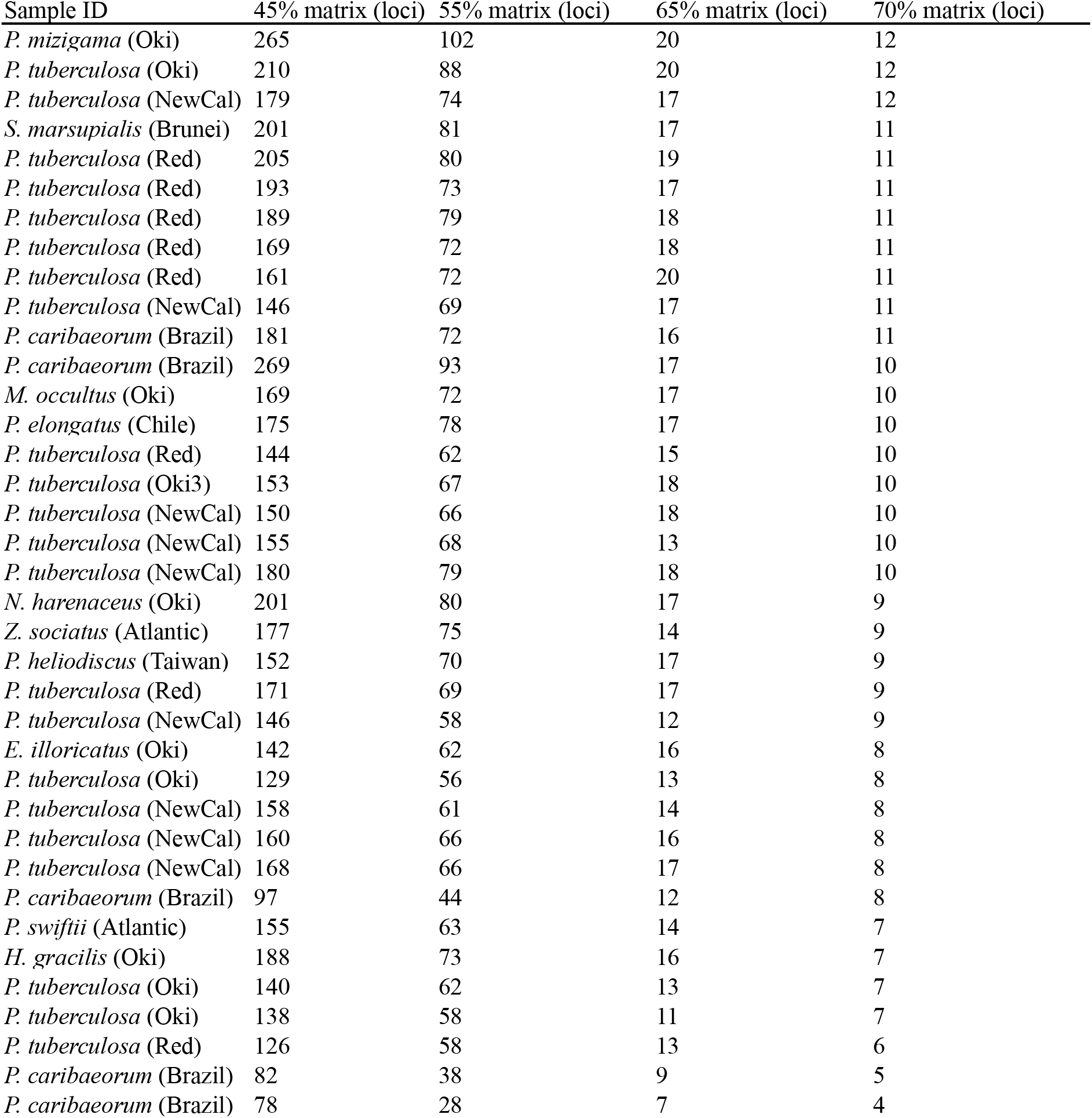
Number of UCE loci recovered per sample across different taxon occupancy thresholds (45%, 55%, 65%, and 70%).

| Sample ID | 45% matrix (loci) | 55% matrix (loci) | 65% matrix (loci) | 70% matrix (loci) |
| --- | --- | --- | --- | --- |
| *P. mizigama* (Oki) | 265 | 102 | 20 | 12 |
| *P. tuberculosa* (Oki) | 210 | 88 | 20 | 12 |
| *P. tuberculosa* (NewCal) | 179 | 74 | 17 | 12 |
| *S. marsupialis* (Brunei) | 201 | 81 | 17 | 11 |
| *P. tuberculosa* (Red) | 205 | 80 | 19 | 11 |
| *P. tuberculosa* (Red) | 193 | 73 | 17 | 11 |
| *P. tuberculosa* (Red) | 189 | 79 | 18 | 11 |
| *P. tuberculosa* (Red) | 169 | 72 | 18 | 11 |
| *P. tuberculosa* (Red) | 161 | 72 | 20 | 11 |
| *P. tuberculosa* (NewCal) | 146 | 69 | 17 | 11 |
| *P. caribaeorum* (Brazil) | 181 | 72 | 16 | 11 |
| *P. caribaeorum* (Brazil) | 269 | 93 | 17 | 10 |
| *M. occultus* (Oki) | 169 | 72 | 17 | 10 |
| *P. elongatus* (Chile) | 175 | 78 | 17 | 10 |
| *P. tuberculosa* (Red) | 144 | 62 | 15 | 10 |
| *P. tuberculosa* (Oki3) | 153 | 67 | 18 | 10 |
| *P. tuberculosa* (NewCal) | 150 | 66 | 18 | 10 |
| *P. tuberculosa* (NewCal) | 155 | 68 | 13 | 10 |
| *P. tuberculosa* (NewCal) | 180 | 79 | 18 | 10 |
| *N. harenaceus* (Oki) | 201 | 80 | 17 | 9 |
| *Z. sociatus* (Atlantic) | 177 | 75 | 14 | 9 |
| *P. heliodiscus* (Taiwan) | 152 | 70 | 17 | 9 |
| *P. tuberculosa* (Red) | 171 | 69 | 17 | 9 |
| *P. tuberculosa* (NewCal) | 146 | 58 | 12 | 9 |
| *E. illoricatus* (Oki) | 142 | 62 | 16 | 8 |
| *P. tuberculosa* (Oki) | 129 | 56 | 13 | 8 |
| *P. tuberculosa* (NewCal) | 158 | 61 | 14 | 8 |
| *P. tuberculosa* (NewCal) | 160 | 66 | 16 | 8 |
| *P. tuberculosa* (NewCal) | 168 | 66 | 17 | 8 |
| *P. caribaeorum* (Brazil) | 97 | 44 | 12 | 8 |
| *P. swiftii* (Atlantic) | 155 | 63 | 14 | 7 |
| *H. gracilis* (Oki) | 188 | 73 | 16 | 7 |
| *P. tuberculosa* (Oki) | 140 | 62 | 13 | 7 |
| *P. tuberculosa* (Oki) | 138 | 58 | 11 | 7 |
| *P. tuberculosa* (Red) | 126 | 58 | 13 | 6 |
| *P. caribaeorum* (Brazil) | 82 | 38 | 9 | 5 |
| *P. caribaeorum* (Brazil) | 78 | 28 | 7 | 4 |

**Supplementary Table 2.** Summary statistics (median, mean, and range) for phylogenetic support metrics across inferred trees, including ultrafast bootstrap support (UFboot), gene concordance factors (gCF), site concordance factors (sCF), and local posterior probabilities (LPP) from ASTRAL analyses.

| Metric | Median | Mean | Range |
| --- | --- | --- | --- |
| UFBoot | 74.5 | 73.9 | 38 - 100 |
| gCF | 3.79 | 18.60 | 0 - 83.5 |
| sCF | 40.30 | 43.21 | 27 - 91.2 |
| ASTRAL LPP | 0.60 | 0.66 | 0.20 - 1.00 |

## REFERENCES

Arrigoni R, Maggioni D, Montano S et al (2018) An integrated morpho-molecular approach to delineate species boundaries of Millepora from the Red Sea. Coral Reefs 37:967–984. 10.1007/s00338-018-01739-8

Beal LM, De Ruijter WPM, Biastoch A et al (2011) On the role of the Agulhas system in ocean circulation and climate. Nature 472:429–436. 10.1038/nature09983

Bolger AM, Lohse M, Usadel B (2014) Trimmomatic: a flexible trimmer for Illumina sequence data. Bioinformatics 30:2114–2120. 10.1093/bioinformatics/btu170

Boscolo HK, Silveira FL (2005) Reproductive biology of Palythoa caribaeorum and Protopalythoa variabilis (Cnidaria, Anthozoa, Zoanthidea) from the southeastern coast of Brazil. Braz J Biol 65:29–41. 10.1590/S1519-69842005000100006

Capella-Gutiérrez S, Silla-Martínez JM, Gabaldón T (2009) trimAl: a tool for automated alignment trimming in large-scale phylogenetic analyses. Bioinformatics 25:1972–1973. 10.1093/bioinformatics/btp348

Carreiro-Silva M, Ocaña O, Stanković D et al (2017) Zoantharians (Hexacorallia: Zoantharia) associated with cold-water corals in the Azores Region: new species and associations in the deep sea. Front Mar Sci 4:88. 10.3389/fmars.2017.00088

Chernomor O, von Haeseler A, Minh BQ (2016) Terrace aware data structure for phylogenomic inference from supermatrices. Syst Biol 65:997–1008. 10.1093/sysbio/syw037

Cord I, Araujo GS, Silva FC et al (2025) Biogeography and evolution of reef fishes on tropical Mid-Atlantic Ridge islands. Proc R Soc B Biol Sci 292:20250756. 10.1098/rspb.2025.0756

Costa D, Gomes P, Santos A et al (2012) Morphological plasticity in the reef zoanthid Palythoa caribaeorum as an adaptive strategy. Ann Zool Fenn 48:349–358. 10.5735/086.048.0602

Cowen RK, Sponaugle S (2009) Larval dispersal and marine population connectivity. Annu Rev Mar Sci 1:443–466. 10.1146/annurev.marine.010908.163757

Cowman PF, Quattrini AM, Bridge TCL et al (2020) An enhanced target-enrichment bait set for Hexacorallia provides phylogenomic resolution of the staghorn corals (Acroporidae) and close relatives. Mol Phylogenet Evol 153:106944. 10.1016/j.ympev.2020.106944

Cruz ICS, Meira VH, de Kikuchi RKP, Creed JC (2016) The role of competition in the phase shift to dominance of the zoanthid Palythoa cf. variabilis on coral reefs. Mar Environ Res 115:28–35. 10.1016/j.marenvres.2016.01.008

Degnan JH, Rosenberg NA (2009) Gene tree discordance, phylogenetic inference and the multispecies coalescent. Trends Ecol Evol 24:332–340. 10.1016/j.tree.2009.01.009

Duchassaing de Fonbressin P, Michelotti G (1860) Mémoire sur les coralliaires des Antilles. Mem Acad Sci Turin Ser 2 19:279–365.

Dudoit A, Iacchei M, Coleman RR et al (2018) The little shrimp that could: phylogeography of the circumtropical Stenopus hispidus (Crustacea: Decapoda) reveals divergent Atlantic and Pacific lineages. PeerJ 6:e4409. 10.7717/peerj.4409

Dudoit A, Santos MEA, Reimer JD, Toonen R (2021) Phylogenomics of Palythoa (Hexacorallia: Zoantharia): probing species boundaries in a globally distributed genus. Coral Reefs 41:1–18. 10.1007/s00338-021-02128-4

Erickson KL, Pentico A, Quattrini AM, McFadden CS (2021) New approaches to species delimitation and population structure of anthozoans: two case studies of octocorals using ultraconserved elements and exons. Mol Ecol Resour 21:78–92. 10.1111/1755-0998.13241

Esper EJC (1791) Die Pflanzenthiere in Abbildungen nach der Natur: mit Farben erleuchtet nebst Beschreibungen. In der Raspischen Buchhandlung, Nürnberg

Faircloth BC (2016) PHYLUCE is a software package for the analysis of conserved genomic loci. Bioinformatics 32:786–788. 10.1093/bioinformatics/btv646

Faircloth BC, McCormack JE, Crawford NG et al (2012) Ultraconserved elements anchor thousands of genetic markers spanning multiple evolutionary timescales. Syst Biol 61:717–726. 10.1093/sysbio/sys004

Faircloth BC, Branstetter MG, White ND, Brady SG (2015) Target enrichment of ultraconserved elements from arthropods provides a genomic perspective on relationships among Hymenoptera. Mol Ecol Resour 15:489–501. 10.1111/1755-0998.12328

Floeter SR, Rocha LA, Robertson DR et al (2008) Atlantic reef fish biogeography and evolution. J Biogeogr 35:22–47. 10.1111/j.1365-2699.2007.01790.x

Giarla TC, Esselstyn JA (2015) The challenges of resolving a rapid, recent radiation: empirical and simulated phylogenomics of Philippine shrews. Syst Biol 64:727–740. 10.1093/sysbio/syv029

Gijsbers JC, Englebert N, Prata KE et al (2023) Global phylogenomic assessment of Leptoseris and Agaricia reveals substantial undescribed diversity at mesophotic depths. BMC Biol 21:147. 10.1186/s12915-023-01630-1

Haywick DW, Mueller EM (1997) Sediment retention in encrusting Palythoa spp.—a biological twist to a geological process. Coral Reefs 16:39–46. 10.1007/s003380050057

Heled J, Drummond AJ (2010) Bayesian inference of species trees from multilocus data. Mol Biol Evol 27:570–580. 10.1093/molbev/msp274

Hirose M, Obuchi M, Hirose E, Reimer JD (2011) Timing of spawning and early development of Palythoa tuberculosa (Anthozoa, Zoantharia, Sphenopidae) in Okinawa, Japan. Biol Bull 220:23–31. 10.1086/BBLv220n1p23

Hoang DT, Chernomor O, von Haeseler A et al (2018) UFBoot2: improving the ultrafast bootstrap approximation. Mol Biol Evol 35:518–522. 10.1093/molbev/msx281

Huang D, Meier R, Todd PA, Chou LM (2008) Slow mitochondrial COI sequence evolution at the base of the metazoan tree and its implications for DNA barcoding. J Mol Evol 66:167–174. 10.1007/s00239-008-9069-5

Kalyaanamoorthy S, Minh BQ, Wong TKF et al (2017) ModelFinder: fast model selection for accurate phylogenetic estimates. Nat Methods 14:587–589. 10.1038/nmeth.4285

Katoh K, Standley DM (2013) MAFFT multiple sequence alignment software version 7: improvements in performance and usability. Mol Biol Evol 30:772–780. 10.1093/molbev/mst010

Kitahara MV, Fukami H, Benzoni F, Huang D (2016) The new systematics of Scleractinia: integrating molecular and morphological evidence. In: Goffredo S, Dubinsky Z (eds) The Cnidaria, past, present and future. Springer, Cham, pp 41–59

Kubatko LS, Degnan JH (2007) Inconsistency of phylogenetic estimates from concatenated data under coalescence. Syst Biol 56:17–24. 10.1080/10635150601146041

Lambre ME, López C, Acha-Araico B, Clemente S (2024) Effects of macroalgae and sea urchin grazing pressure on zoantharians growth under laboratory conditions. Mar Environ Res 198:106534. 10.1016/j.marenvres.2024.106534

Letunic I, Bork P (2024) Interactive Tree of Life (iTOL) v6: recent updates to the phylogenetic tree display and annotation tool. Nucleic Acids Res 52:W78–W82. 10.1093/nar/gkae268

López C, Reimer JD, Brito A et al (2019) Diversity of zoantharian species and their symbionts from the Macaronesian and Cape Verde ecoregions demonstrates their widespread distribution in the Atlantic Ocean. Coral Reefs 38:269–283. 10.1007/s00338-019-01773-0

Low M, Sinniger F, Reimer J (2016) The order Zoantharia Rafinesque, 1815 (Cnidaria: Anthozoa: Hexacorallia): supraspecific classification and nomenclature. ZooKeys 641:1–49. 10.3897/zookeys.641.10346

Lustic C, Maxwell K, Bartels E et al (2020) The impacts of competitive interactions on coral colonies after transplantation: a multispecies experiment from the Florida Keys, US. Bull Mar Sci 96:805–818. 10.5343/bms.2019.0086

Maddison WP (1997) Gene trees in species trees. Syst Biol 46:523–536

McFadden CS, Quattrini AM, Brugler MR et al (2021) Phylogenomics, origin, and diversification of Anthozoans (Phylum Cnidaria). Syst Biol 70:635–647. 10.1093/sysbio/syaa103

McFadden CS, Erickson KL, Lane A et al (2025) Biodiversity and biogeography of zooxanthellate soft corals across the Indo-Pacific. Sci Rep 15:15461. 10.1038/s41598-025-98790-7

Minh BQ, Schmidt HA, Chernomor O et al (2020) IQ-TREE 2: new models and efficient methods for phylogenetic inference in the genomic era. Mol Biol Evol 37:1530–1534. 10.1093/molbev/msaa015

Mirarab S, Reaz R, Bayzid MdS et al (2014) ASTRAL: genome-scale coalescent-based species tree estimation. Bioinformatics 30:i541–i548. 10.1093/bioinformatics/btu462

Mizuyama M, Masucci GD, Reimer JD (2018) Speciation among sympatric lineages in the genus Palythoa (Cnidaria: Anthozoa: Zoantharia) revealed by morphological comparison, phylogenetic analyses and investigation of spawning period. PeerJ 6:e5132. 10.7717/peerj.5132

Moreno-Borges S, López C, Clemente S (2022) Reef fish assemblages associated to new mat-forming zoantharian communities in the Canary Islands. Mar Environ Res 177:105623. 10.1016/j.marenvres.2022.105623

Nguyen L-T, Schmidt HA, von Haeseler A, Minh BQ (2015) IQ-TREE: a fast and effective stochastic algorithm for estimating maximum-likelihood phylogenies. Mol Biol Evol 32:268–274. 10.1093/molbev/msu300

Ong CW, Reimer JD, Todd PA (2013) Morphologically plastic responses to shading in the zoanthids Zoanthus sansibaricus and Palythoa tuberculosa. Mar Biol 160:1053–1064. 10.1007/s00227-012-2158-4

Oury N, Noël C, Mona S et al (2023) From genomics to integrative species delimitation? The case study of the Indo-Pacific Pocillopora corals. Mol Phylogenet Evol 184:107803. 10.1016/j.ympev.2023.107803

Polak O, Loya Y, Brickner I et al (2011) The widely-distributed Indo-Pacific zoanthid Palythoa tuberculosa: a sexually conservative strategist. Bull Mar Sci 87:605–621. 10.5343/bms.2010.1088

Prjibelski A, Antipov D, Meleshko D et al (2020) Using SPAdes de novo assembler. Curr Protoc Bioinformatics 70:e102. 10.1002/cpbi.102

Quattrini AM, Faircloth BC, Dueñas LF et al (2018) Universal target-enrichment baits for anthozoan (Cnidaria) phylogenomics: new approaches to long-standing problems. Mol Ecol Resour 18:281–295. 10.1111/1755-0998.12736

Quattrini AM, Snyder KE, Purow-Ruderman R et al (2023) Mito-nuclear discordance within Anthozoa, with notes on unique properties of their mitochondrial genomes. Sci Rep 13:7443. 10.1038/s41598-023-34059-1

Reimer JD, Fujii T (2010) Four new species and one new genus of zoanthids (Cnidaria, Hexacorallia) from the Galapagos Islands. ZooKeys 42:1–36. 10.3897/zookeys.42.378

Reimer JD, Ono S, Takishita K et al (2006) Molecular evidence suggesting species in the zoanthid genera Palythoa and Protopalythoa (Anthozoa: Hexacorallia) are congeneric. Zoolog Sci 23:87–94. 10.2108/zsj.23.87

Reimer JD, Takishita K, Ono S, Maruyama T (2007) Diversity and evolution in the zoanthid genus Palythoa (Cnidaria: Hexacorallia) based on nuclear ITS-rDNA. Coral Reefs 26:399–410. 10.1007/s00338-007-0210-5

Reimer JD, Nakachi S, Hirose M et al (2010) Using hydrofluoric acid for morphological investigations of zoanthids (Cnidaria: Anthozoa): a critical assessment of methodology and necessity. Mar Biotechnol 12:605–617. 10.1007/s10126-009-9249-3

Reimer JD, Wee HB, López C et al (2021) Widespread Zoanthus and Palythoa dominance, barrens, and phase shifts in shallow water subtropical and tropical marine ecosystems. In: Hawkins SJ et al (eds) Oceanography and Marine Biology: An Annual Review. CRC Press, Boca Raton, pp 533–557

Rieux A, Balloux F (2016) Inferences from tip-calibrated phylogenies: a review and a practical guide. Mol Ecol 25:1911–1924. 10.1111/mec.13586

Rosser NL, Thomas L, Stankowski S et al (2017) Phylogenomics provides new insight into evolutionary relationships and genealogical discordance in the reef-building coral genus Acropora. Proc R Soc B Biol Sci 284:20162182. 10.1098/rspb.2016.2182

Ruiz-Ramos DV, Weil E, Schizas NV (2014) Morphological and genetic evaluation of the hydrocoral Millepora species complex in the Caribbean. Zool Stud 53:4. 10.1186/1810-522X-53-4

Ryland JS, Putron SD, Scheltema RS et al (2000) Semper’s (zoanthid) larvae: pelagic life, parentage and other problems. Hydrobiologia 440:191–198. 10.1023/A:1004171127777

Santos MEA, Reimer JD (2018) Rafting in Zoantharia: a hitchhiker’s guide to dispersal? Mar Pollut Bull 130:307–310. 10.1016/j.marpolbul.2018.03.041

Santos MEA, Reimer JD, Kiriukhin B et al (2024) Coral microbiomes from the Atlantic and Indo-Pacific oceans have the same alpha diversity but different composition. bioRxiv (preprint) 10.1101/2024.08.16.608269

Santos MEA, Kise H, Fourreau CJL et al (2026) Global biogeography of zoantharians indicates a weak genetic differentiation between the Atlantic and Indo-Pacific oceans, and distinct communities in tropical and temperate provinces. Front Biogeogr 19:e174247. 10.21425/fob.19.174247

Schleyer MH, Floros C, Laing SCS et al (2018) What can South African reefs tell us about the future of high-latitude coral systems? Mar Pollut Bull 136:491–507. 10.1016/j.marpolbul.2018.09.014

Schwaninger H (2008) Global mitochondrial DNA phylogeography and biogeographic history of the antitropically and longitudinally disjunct marine bryozoan Membranipora membranacea L. (Cheilostomata): Another cryptic marine sibling species complex? Mol Phylogenet Evol 49:893–908. 10.1016/j.ympev.2008.08.016

Shearer TL, van Oppen MJH, Romano SL, Wörheide G (2002) Slow mitochondrial DNA sequence evolution in the Anthozoa (Cnidaria). Mol Ecol 11:2475–2487. 10.1046/j.1365-294X.2002.01652.x

Sinniger F, Montoya-Burgos J, Chevaldonné P, Pawlowski J (2005) Phylogeny of the order Zoantharia (Anthozoa, Hexacorallia) based on the mitochondrial ribosomal genes. Mar Biol 147:1121–1128. 10.1007/s00227-005-0016-3

Sinniger F, Reimer JD, Pawlowski J (2010) The Parazoanthidae (Hexacorallia: Zoantharia) DNA taxonomy: description of two new genera. Mar Biodivers 40:57–70. 10.1007/s12526-009-0034-3

Stelbrink B, von Rintelen T, Cliff G, Kriwet J (2010) Molecular systematics and global phylogeography of angel sharks (genus Squatina). Mol Phylogenet Evol 54:395–404. 10.1016/j.ympev.2009.07.029

Tan X, Qi J, Liu Z et al (2023) Phylogenomics reveals high levels of incomplete lineage sorting at the ancestral nodes of the macaque radiation. Mol Biol Evol 40:msad229. 10.1093/molbev/msad229

Veron J, Stafford-Smith M, DeVantier L, Turak E (2015) Overview of distribution patterns of zooxanthellate Scleractinia. Front Mar Sci 1:81. 10.3389/fmars.2014.00081

Whitfield JB, Lockhart PJ (2007) Deciphering ancient rapid radiations. Trends Ecol Evol 22:258–265. 10.1016/j.tree.2007.01.012

Zamora-Jordán N, Martínez P, Hernández M, López C (2024) Responses of Palythoa caribaeorum and its associated endosymbionts to thermal stress. Coral Reefs 43:1443–1454. 10.1007/s00338-024-02549-x

Zamora-Jordán N, Hernández M, Reimer JD et al (2026) Molecular analyses of the Palythoa caribaeorum holobiont suggest its recent arrival in the Canary Islands. Mar Ecol Prog Ser 779:1–15. 10.3354/meps15060

Zhang C, Rabiee M, Sayyari E, Mirarab S (2018) ASTRAL-III: polynomial time species tree reconstruction from partially resolved gene trees. BMC Bioinformatics 19:153. 10.1186/s12859-018-2129-y

